# Voluntary and forced exposure to ethanol vapor produces similar escalation of alcohol drinking but differential recruitment of brain regions related to stress, habit, and reward in male rats

**DOI:** 10.1101/2022.05.11.491572

**Authors:** Giordano de Guglielmo, Sierra Simpson, Adam Kimbrough, Dana Conlisk, Robert Baker, Maxwell Cantor, Marsida Kallupi, Olivier George

**Author notes:** <u>Corresponding author:</u> Giordano de Guglielmo, Department of Psychiatry, University of California San Diego, 9500 Gilman Drive, 92093 La Jolla, CA, USA.

## Abstract

A major limitation of the most widely used current animal models of alcohol dependence is that they use forced exposure to ethanol including ethanol-containing liquid diet and chronic intermittent ethanol (CIE) vapor to produce clinically relevant blood alcohol levels (BAL) and addiction-like behaviors. We recently developed a novel animal model of voluntary induction of alcohol dependence using ethanol vapor self-administration (EVSA). In this model, naive outbred rats given intermittent access to alcohol vapor self-administration exhibit BAL in the 150-300 mg% range and develop somatic signs of withdrawal during acute abstinence. However, it is unknown whether EVSA leads to an escalation of alcohol drinking *per se*, and whether such escalation is associated with neuroadaptations in brain regions related to stress, reward, and habit. To address these issues, we compared the levels of alcohol drinking during withdrawal between rats passively exposed to alcohol (CIE) or voluntarily exposed to EVSA and measured the number of Fos+ neurons during acute withdrawal (16 h) in the central nucleus of the amygdala (CeA), dorsomedial striatum (DMS), dorsolateral striatum (DLS), nucleus accumbens core (Nacc), periaqueducal grey area (PAG), lateral Habenula (HbL), and the paraventricular nucleus of the thalamus (PVT). The rats were first trained to orally self-administer alcohol in standard operant chambers and then divided in 4 groups (CIE, CI-Air, EVSA and Air-SA) and exposed to intermittent ethanol vapor (passive or active) or intermittent air (passive or active) for 8 h/day, 3 days a week. CIE and EVSA rats exhibited similar BAL (150-300 mg% range) and similar escalation of alcohol drinking during withdrawal, while no changes in terms of drinking were observed in the air exposed rats. CIE and EVSA also increased the motivation for alcohol compared to their respective air control groups under a progressive ratio schedule of reinforcement. Acute withdrawal from EVSA and CIE recruited a similar number of Fos+ neurons in the CeA, however, acute withdrawal from EVSA recruited a higher number of Fos+ neurons in every other brain region analyzed compared to acute withdrawal from CIE. Moreover, acute withdrawal from EVSA specifically recruited the DMS and PVT, a pattern not observed in CIE rats.

In summary, these results demonstrate that EVSA produces similar escalation of alcohol drinking, motivation to drink, and blood-alcohol levels than the CIE model, while letting animals voluntary initiate alcohol exposure and maintain alcohol dependence. Moreover, while the behavioral measures of alcohol dependence between the voluntary (EVSA) and passive (CIE) model was similar, the recruitment of neuronal ensembles during acute withdrawal was very different with a higher recruitment of Fos+ neurons in key brain regions important for stress, reward and habit-related processes. The EVSA model may be particularly useful to unveil the neuronal networks and pharmacology responsible for the voluntary induction and maintenance of alcohol dependence and may improve translational studies by providing preclinical researchers with an animal model with better face validity for alcohol use disorder.

## Introduction

A major problem in the alcohol field is the limited number of animal models of alcohol self-administration in outbred rats that leads to blood alcohol levels in the 150-300 mg% range, a level often observed in alcohol use disorder and extreme binge drinking. The lack of animal model hinders the discovery of the neuronal networks responsible for the voluntary initiation and maintenance of alcohol use disorder and the discovery of novel pharmacological treatments.

Although no animal model of addiction fully emulates the human condition, the models do permit the investigation of the specific elements of the addiction process [1]. Multiple methods have been developed to produce BALs in the 100-300 mg% range for 12+ h per day, associated with somatic and affective withdrawal symptoms, but they either use forced alcohol liquid diet [2–4], intragastric ethanol intubation [5–8], passive exposure to ethanol vapor [9–15], or selected inbred lines of ethanol-preferring rats or mice [16,17].

The current gold standard animal model for the induction of alcohol dependence in rats is the model of chronic intermittent ethanol vapor (CIE). The CIE model represents a very efficient way to produce alcohol dependence [3,15,18–22], and over the years it has been extremely helpful to elucidate the neuronal mechanisms underlying alcohol dependence [18,19,23–38].

We recently characterized a novel animal model animal for the induction and maintenance of alcohol dependence in outbred rats using chronic intermittent alcohol vapor self-administration (EVSA model [39]) by using a novel apparatus that allow rats to self-administer alcohol vapor in their home cage. Using this innovative apparatus and a new intermittent access paradigm, we found that rats self-administer ethanol vapor to the point of reaching BALs in the 150-300 mg% range and show an addicted-like phenotype (escalation of intake anxiety-like behavior, hyperalgesia, somatic withdrawal signs) similar to passive ethanol vapor exposure using the CIE model [39].

However, it is unknown whether EVSA leads to an escalation of alcohol drinking *per se*,and whether such escalation is associated with neuroadaptations in brain regions important for stress, reward, and habit-related processes. The goal of this study was to compare the levels of alcohol drinking and motivation for alcohol during withdrawal in animals passively exposed to alcohol (CIE) and animals subjected to the new ethanol vapor self-administration paradigm (EVSA). We also performed neuroanatomical analysis of the immediate early gene *c-fos* in several addiction-related brain regions and compared patterns of activation between CIE and EVSA rats during acute alcohol withdrawal (16 h).

## Materials and Methods

A schematic of the experimental design is represented in Figure 1.

**Figure 1:**
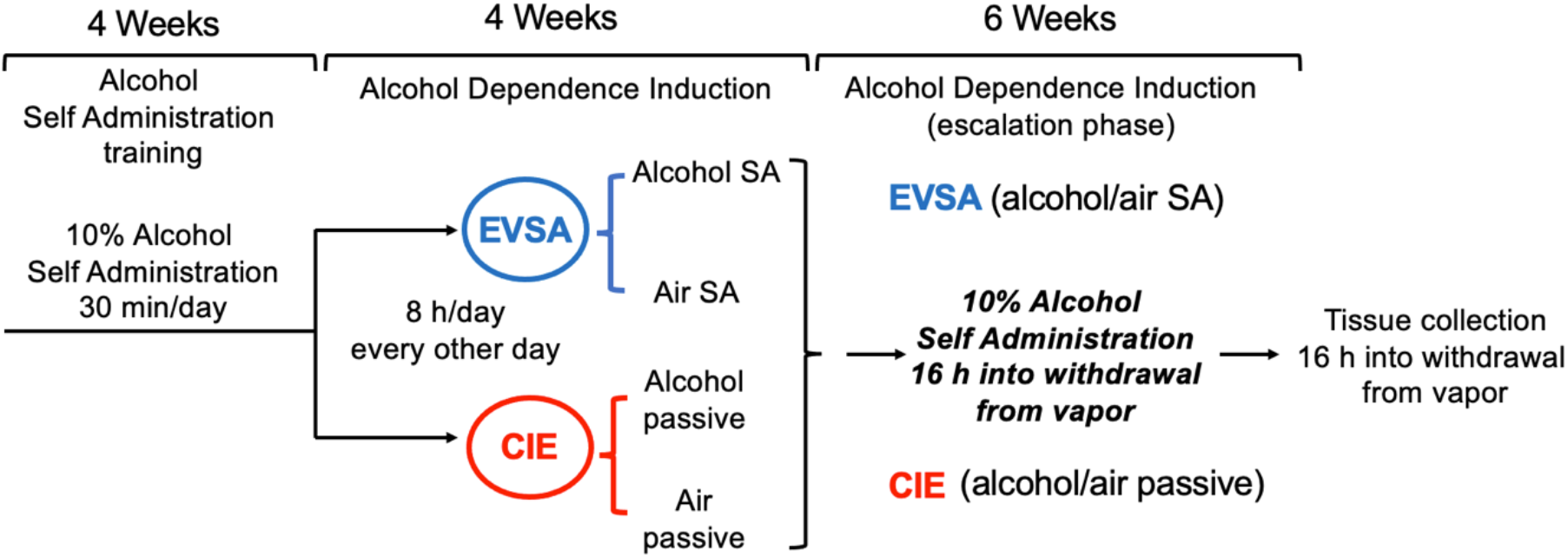
Experimental Timeline. The rats were first trained to self-administer alcohol in standard operant chambers and then divided in 4 groups (CIE, Air Passive, EVSA and Air SA, and exposed to intermittent ethanol vapor (passive or active) or intermittent air (control groups passive or active) 8 h/day 3 days a week for 4 weeks. The animals were then tested for alcohol drinking in the operant chambers 16 hours into acute withdrawal from vapor for 6 more weeks.

### Subjects

Adult male *Wistar* rats (n=40), weighing 200-225 g at the beginning of the experiments, were housed in groups of two per cage (self-administration groups) in a temperature-controlled (22°C) vivarium on a 12 h/12 h light/dark cycle (lights on at 10:00 PM) with *ad libitum* access to food and water. All the behavioral tests were conducted during the dark phase of the light/dark cycle. All the procedures adhered to the National Institutes of Health Guide for the Care and Use of Laboratory Animals and were approved by the Institutional Animal Care and Use Committee of the University of California, San Diego.

### Operant self-administration (drinking)

Self-administration sessions were conducted in standard operant conditioning chambers (Med Associates, St. Albans, VT, USA). For the alcohol self-administration studies, the animals were first trained to self-administer 10% (w/v) alcohol and water solutions until a stable response pattern (20 ± 5 rewards) was maintained. The rats (n=32) were subjected to an overnight session in the operant chambers with access to one lever (right lever) that delivered water (fixed-ratio 1 [FR1]). Food was available *ad libitum* during this training. After 1 day off, the rats were subjected to a 2 h session (FR1) for 1 day and a 1 h session (FR1) the next day, with one lever delivering alcohol (right lever). All the subsequent alcohol self-administration sessions lasted 30 min. The rats were allowed to self-administer a 10% (w/v) alcohol solution (right lever) and water (left lever) on an FR1 schedule of reinforcement (i.e., each operant response was reinforced with 0.1 ml of the solution). Lever presses on the left lever delivered 0.1 ml of water. This procedure lasted 12 days until a stable baseline of intake was reached. At the end of this phase the rats were divided into 4 groups (n=8/group) that underwent the EVSA procedure (alcohol or air) or the CIE procedure (alcohol or air). A schematic of the experiment is presented in Figure 1. Eight more age-matched naïve animals were used at the end of the study as a naïve control for the FOS expression experiment.

### Operant ethanol vapor self-administration (EVSA)

We recently developed a new apparatus that consists of a standard rat home cage that is equipped with a nosepoke hole and a cue light on each side of the chamber for active and inactive responses [39]. The apparatus was connected to a custom-made alcohol vaporization system, consisting of a heating element, a glass flask, two solenoids (one “normally opened,” connected to clean air; one “normally closed,” connected to the alcohol flask), a gas washing bottle, and a compressor. The apparatus was controlled by a Med Associates smartcard, and each response at the active nosepoke hole triggered the activation of the two solenoids. The one that was “normally open” closed, and the one that was “normally closed” opened, so alcohol vapor could be released into the chamber. After 2, 5, or 10 min (depending on the parameters used), the solenoids turned off, and clean air was pushed into the chamber. All active and inactive responses were recorded on a computer. The home cage was always ventilated with a minimum of 15 L/min of clean air during the experiment.

### Chronic-intermittent alcohol/air vapor self-administration model (EVSA)

The animals (n=8) were trained to self-administer alcohol vapor in 8 h sessions every other day from 10 AM to 6 PM for a total of 12 sessions. Rats did not have access to food or water during the test sessions. For the first 4 sessions, a response in the active nosepoke hole resulted in the activation of two solenoids (one for alcohol vapor opened and one for clean air closed) to expose the animals to alcohol vapor (15 L/min) for 2 consecutive minutes. The illumination of a cue light for 20 s followed a nosepoke response. Any nosepoke in the active hole while the cue light was on did not result in any delivery of alcohol vapor (timeout [TO]). After the 20 s TO, responses in the active nosepoke hole released alcohol vapor or air for another 2 min, and the cue light was illuminated again for other 20 s. Responses in the opposite hole were recorded but had no consequences. In the next four sessions, the animals were subjected to the same parameters. The only difference was that the time of vapor exposure after each nosepoke increased to 5 min. In the last four sessions, the time of vapor exposure after each nosepoke increased to 10 min. A separate group of 8 rats was used as a control and was trained in the same conditions with the only difference that the response to the active nose poke triggered the release of air instead of alcohol (Figure 1).

### Alcohol vapor chambers (Chronic Intermittent Ethanol vapor exposure, CIE, model)

A separate group of rats (n=8) was made dependent by chronic intermittent exposure to alcohol vapors. They underwent cycles of 8 h every other day (blood alcohol levels during vapor exposure ranged between 150 and 250 mg%) for 4 weeks. The every other day 8h exposure has been chosen to match the 8 h paradigm in the EVSA model. In this model, rats exhibit somatic and motivational signs of withdrawal [14]. Eight more rats were used as a control and exposed to air (Figure 1).

### Operant self-administration during alcohol vapor (active or passive) exposure

Behavioral testing occurred three times per week 16 ours into withdrawal from alcohol/air vapor. The rats were tested for alcohol (and water) self-administration on an FR1 schedule of reinforcement in 30-min sessions for 6 weeks. At the end of this phase the animals were subjected to a progressive ratio test during which the number of lever presses necessary to obtain the next reinforcement progressively increased according to the following progression: 1, 1, 2, 2, 3, 3, 4, 4, 5, 5, 7, 7, 9, 9, 11, 11, 13, 13, etc. [19,26]. The PR session stopped after 1.30 h or when 30 min elapsed after the last reinforcement.

### Determination of blood alcohol levels

Blood was sampled weekly for the determination of BALs during vapor exposure for both the EVSA and the CIE paradigms. The rats were gently restrained under the technician’s arm, and the tip of the tail (2 mm) was cut with a clean razor blade.

Blood samples (0.1 ml) were collected in Eppendorf tubes (catalog no. 05-407-13C, Fisher Scientific, Waltham, MA) that contained evaporated heparin and kept on ice. The samples were centrifuged, and serum was decanted into fresh Eppendorf tubes. The serum was then processed using GAS Chromatography for BAL determination.

### Immunohistochemistry

At the end of the experiments, the rats were deeply anesthetized with C02 16 h into withdrawal at the time they were normally tested for alcohol drinking and perfused with 100 ml of phosphate-buffered saline (PBS) followed by 400 ml of 4% paraformaldehyde. The brains were postfixed in paraformaldehyde overnight and transferred to 30% sucrose in PBS/0.1% azide solution at 4°C for 2-3 days.

Brains were frozen in powdered dry ice and sectioned on a cryostat. Coronal sections were cut 40 μm thick and collected free-floating in PBS/0.1% azide solution. Following three washes in PBS, the sections were incubated in 1% hydrogen peroxide/PBS (to quench endogenous peroxidase activity), rinsed three times in PBS, and blocked for 60 min in PBS that contained 0.3% TritonX-100, 1 mg/ml bovine serum albumin, and 5% normal donkey serum. Sections were incubated for 24 h at 4°C with a rabbit monoclonal anti-Fos antibody (Cell Signaling Technologies #2250) diluted 1:1000 in PBS/0.5%Tween20 and 5% normal donkey serum. The sections were washed again with PBS and incubated for 1 h in undiluted Rabbit ImmPress HRP reagent (Vector Laboratories). After washing in PBS, the sections were developed for 2-6 min in Vector peroxidase DAB substrate (Vector Labs) enhanced with nickel chloride. Following PBS rinses, the sections were mounted onto coated slides (Fisher Super Frost Plus), air dried, dehydrated through a graded series of alcohol, cleared with Citrasolv (Fisher Scientific), and coverslipped with DPX (Sigma).

Quantitative analysis to obtain unbiased estimates of the total number of Fos+ cell bodies was performed on a Zeiss Axiophot Microscope equipped with MicroBrightField Stereo Investigator software (Colchester, VT, USA), a three-axis Mac 5000 motorized stage (Ludl Electronics Products, Hawthorne, NY, USA), a Q Imaging Retiga 2000R color digital camera, and PCI color frame grabber.

Three sections were bilaterally analyzed for each rat. Live video images were used to draw contours (at 5 x magnification) to delineate the DMS, DLS, NAcc, CeA, PAG, Medial Habenula and PVT. After determining the mounted section thickness and Z-plane values, an optical fractionator probe was used to determine the number of positive neurons within the DMS, DLS, NAcc, CeA, PAG, Medial Habenula and PVT contours. Briefly, a counting frame of appropriate dimensions, denoting forbidden and non-forbidden boundaries, was superimposed on the video monitor, and Fos+ cells were counted at 20× magnification. Cells were identified as neurons based on standard morphology, and only neurons with a focused nucleus within the nonforbidden regions of the counting frame were counted. Counts from all images from each rat were averaged so that each rat was an *n* of 1.

### Statistical analysis

The data are expressed as mean ± SEM. For comparisons between only two groups, *p* values were calculated using unpaired *t*-tests as described in the result section. Comparisons across more than two groups were made using one-way analysis of variance (ANOVA), and two-way ANOVA was used when there was more than one independent variable. The standard error of the mean is indicated by error bars for each group of data. Differences were considered significant at *p* < 0.05. All of these data were analyzed using Statistica 7 software.

## Results

### Pre-dependence alcohol self-administration

The animals successfully learned to self-administer alcohol and to discriminate between alcohol and water during the 12 days of the training phase, as demonstrated by the significant interaction in Two Way ANOVA, with solutions (Alcohol/Water) as between factor and time (number of sessions) as within factor (F_9,558_ = 3.459, *p* < 0.001). The Newman Keuls post hoc test showed that animals significantly preferred alcohol over water in the last 3 days of self-administration (*p* < 0.01, Fig. 2A). At this point the animals were divided into 4 groups (EVSA alcohol, EVSA air, CIE alcohol, CIE air) with similar alcohol and water intake as shown by the non-significant 2 Way ANOVAs (F_27,252_ = 0.97, *p* = NS, for alcohol, Fig. 2B and F_27,252_ = 0.70, *p* = NS, for water, Fig. 2C).

**Figure 2:**
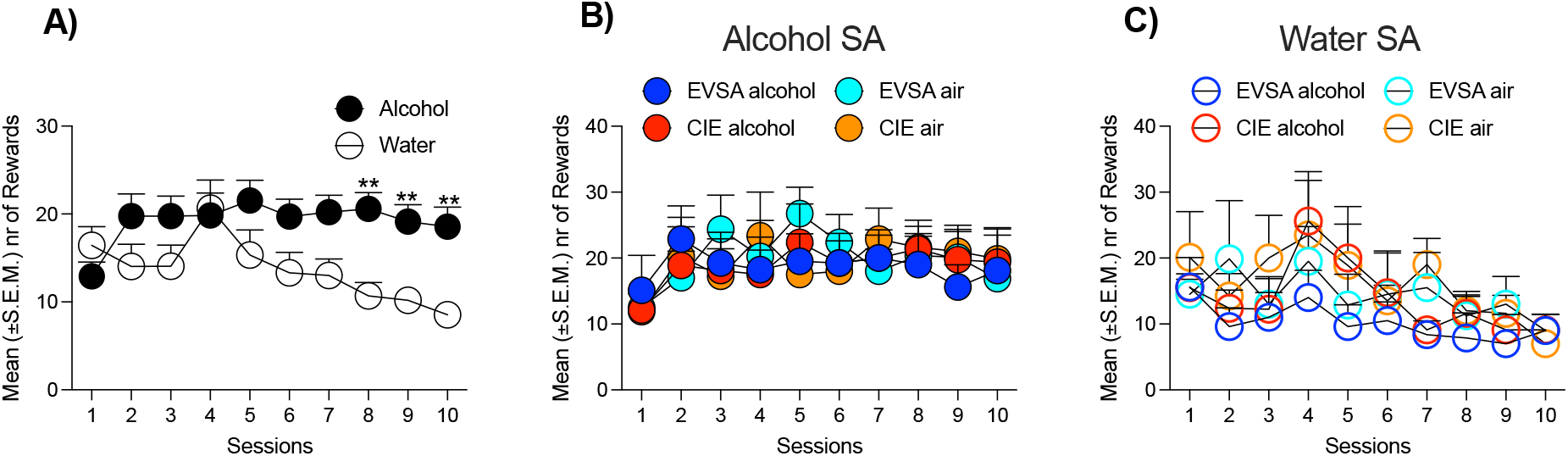
Alcohol Self Administration pre-vapor exposure. **A**) Discrimination between alcohol and water levers in the whole population **B**) Mean of alcohol rewards in the 4 groups before alcohol vapor/air exposure **C**) Mean of water rewards in the 4 groups before alcohol vapor/air exposure. ** *p* < 0.01 vs water

### Ethanol vapor self-administration (EVSA)

At this point the CIE rats were placed in vapor chambers for 3 weeks, while the EVSA rats started self-administering alcohol vapor or air in 8 hours daily sessions for 3 weeks.

The rats self-administering alcohol vapor discriminated very well between the active and inactive nosepoke operanda (Fig. 3A). The mixed factorial ANOVA, with nose pokes as the between-subjects factor and session as the within-subjects factor, revealed a significant nose pokes × session interaction (*F*_11,154_ = 2.918, *p* < 0.001). In the first session, the animals emitted 64.9% (102.1 ± 14.6) active nosepokes *vs*. 35.1% (54.2 ± 7.4) inactive nosepokes. The percentage of active *vs*. inactive nosepokes increased as the number of sessions increased. In the last four sessions, the animals exhibited strong preference for the active nosepoke hole (80.5%, 47.6 ± 13.4 *vs*. 8.2 ± 2.2).

**Figure 3:**
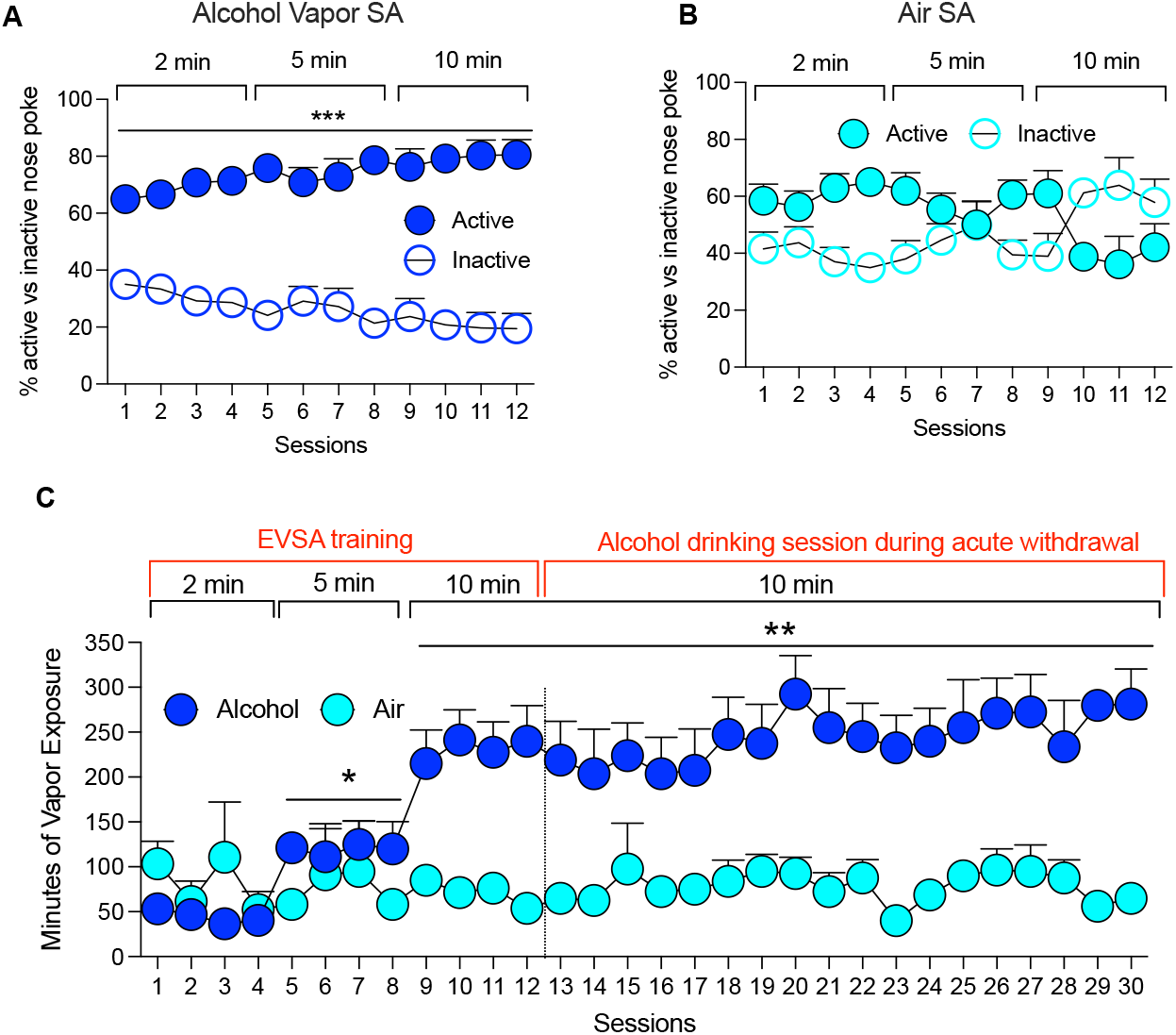
Alcohol/air vapor self-administration. **A**) Discrimination between active and inactive responses in the EVSA group, \*\*\**p* < 0.01 vs inactive **B**) Discrimination between active and inactive responses in the air self-administration group. **C**) The rats nose-poking for alcohol escalate their intake while the rats nose-poking for air show relatively low levels of responding \*\*\**p* < 0.05 and ** *p* < 0.01 vs last day of 2 min phase.

On the other hand, the control rats self-administering air did not show discrimination between the active and inactive nosepoke operanda (Fig. 3B). While in the first few sessions the rats seemed to prefer the active nose poke (associated with the activation of the cue light), their preference shifted for the inactive one in the last few sessions of the training paradigm (Fig. 3B). The animals self-administering alcohol vapor showed robust escalation of alcohol vapor self-administration as demonstrated by the significant drug x time interaction in the two-way ANOVA (*F*_29,406_ = 6.063, *p* < 0.0001) that was not observed with the rats self-administering air. The Newman Keuls *post hoc* test revealed that alcohol animals escalated their exposure to alcohol vapor from session 5 to session 8 (*p* < 0.05, 5 min phase) and from session 9 to session 30 (*p* < 0.01, 10 min phase; Fig 3C) compared to session 4 (last session at 2 min vapor reward).

Air self-administration rates in the air group remained low for the whole duration of the experiment.

### Escalation of alcohol drinking during withdrawal

After 3 weeks of passive (CIE) or active (EVSA) alcohol exposure there was no difference in blood alcohol level concentrations between CIE and EVSA rats as demonstrated by the nonsignificant t-test (*t*_14_ = 0.14, *p* = NS; Fig. 4A). The animals were maintained in the vapor (CIE) or let self-administer alcohol vapor (EVSA) for 6 more weeks and tested 18 hours into withdrawal from vapor on 18 30 min alcohol self-administration sessions. The two-way ANOVA with drug as between factor and time as within factor showed a significant drug x time interaction (*F*_60,560_ = 2.389, *p* < 0.0001). Analysis of the Newman Keuls post hoc demonstrated that CIE and EVSA rats escalated their alcohol compared to their pre-dependence baseline, while no escalation was detected in their respective air control animals (Fig 4B). Figure 4C shows comparisons of the intake pre- and post-vapor exposure in the four groups. The two-way ANOVA with drug as between factor and time as within factor showed a significant drug x time interaction (*F*_3,28_ = 15.45, *p* < 0.0001). The Newman Keuls *post hoc* test revealed that both CIE and EVSA rats escalated their alcohol intake (*p* < 0.001) compared to their pre-dependence baseline and consumed significant more alcohol compared to their air counterpart after alcohol vapor exposure (*p* < 0.001). No difference in terms of alcohol intake was detected between EVSA and CIE, demonstrating that the two models of ethanol dependence induction led to similar alcohol intake during the escalation phase.

**Figure 4:**
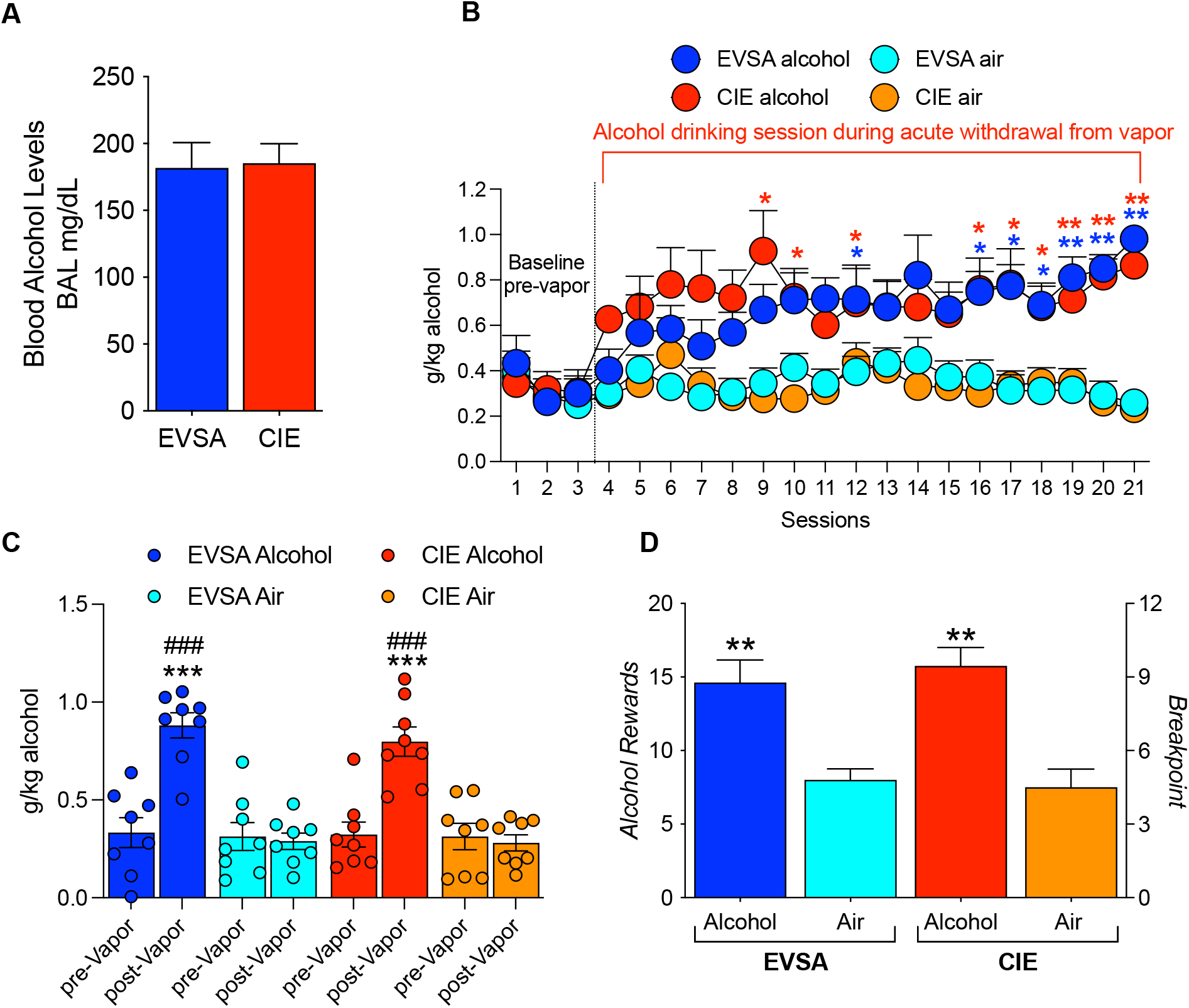
CIE and EVSA alcohol rats show similar BALs and similar level of escalation and motivation for alcohol. **A**) Blood alcohol levels (BALs) at the end of the vapor exposure. **B**) Escalation of alcohol drinking in the 4 groups. CIE and EVSA alcohol rats show escalation of alcohol intake when subjected to a drinking session during acute withdrawal. \**p* < 0.05 and ** *p* < 0.01 vs Baseline pre vapor. **C**) Average for the levels of alcohol drinking pre-(last 3 days) and post (last 3 days) vapor exposure. \*\*\**p* < 0.001 vs Baseline pre vapor and ###*p* < 0.001 vs air control. (**D**) Motivation for alcohol, measured in a Progressive Ratio test, at the end of the behavioral paradigm. ** *p* < 0.01 vs air control.

The rats were also tested in a progressive ratio paradigm. The two-way ANOVA with drug as between factor and time as within factor showed a significant drug x time interaction (*F*_3,28_ = 15.45, *p* < 0.0001). The Newman Keuls *post hoc* test revealed that both CIE and EVSA alcohol rats showed increased motivation for alcohol compared their air exposed control groups (*p* < 0.001, Fig.4D). No difference in the progressive ratio test was detected between EVSA and CIE, demonstrating that the two models of ethanol dependence induction led to similar motivation for alcohol during acute withdrawal. At the end of the behavioral studies, brains were collected for the evaluation of Fos+ cells in different brain regions

### Characterization of neuronal ensembles recruited by EVSA and CIE during acute withdrawal from alcohol vapor

We then investigated the neuronal ensembles that are recruited by acute withdrawal from alcohol vapor in animals made dependent through EVSA and CIE by evaluating the number of Fos-positive neurons in key brain regions that are involved in addiction (DMS, DLS, NAcc, CeA, PAG, Medial Habenula and PVT). The animals were euthanized 16 hours into withdrawal at the time they were used to start the alcohol self-administration testing. Eight more age-matched naïve animals were euthanized at the same time point and used as naïve controls.

Figure 5 depicts the number of Fos-positive neurons per millimeter squared in the DMS, DLS, NAcc, CeA, PAG, Medial Habenula and PVT. Each graph shows the mean ± SEM (n = 6–8 animals per group).

**Figure 5:**
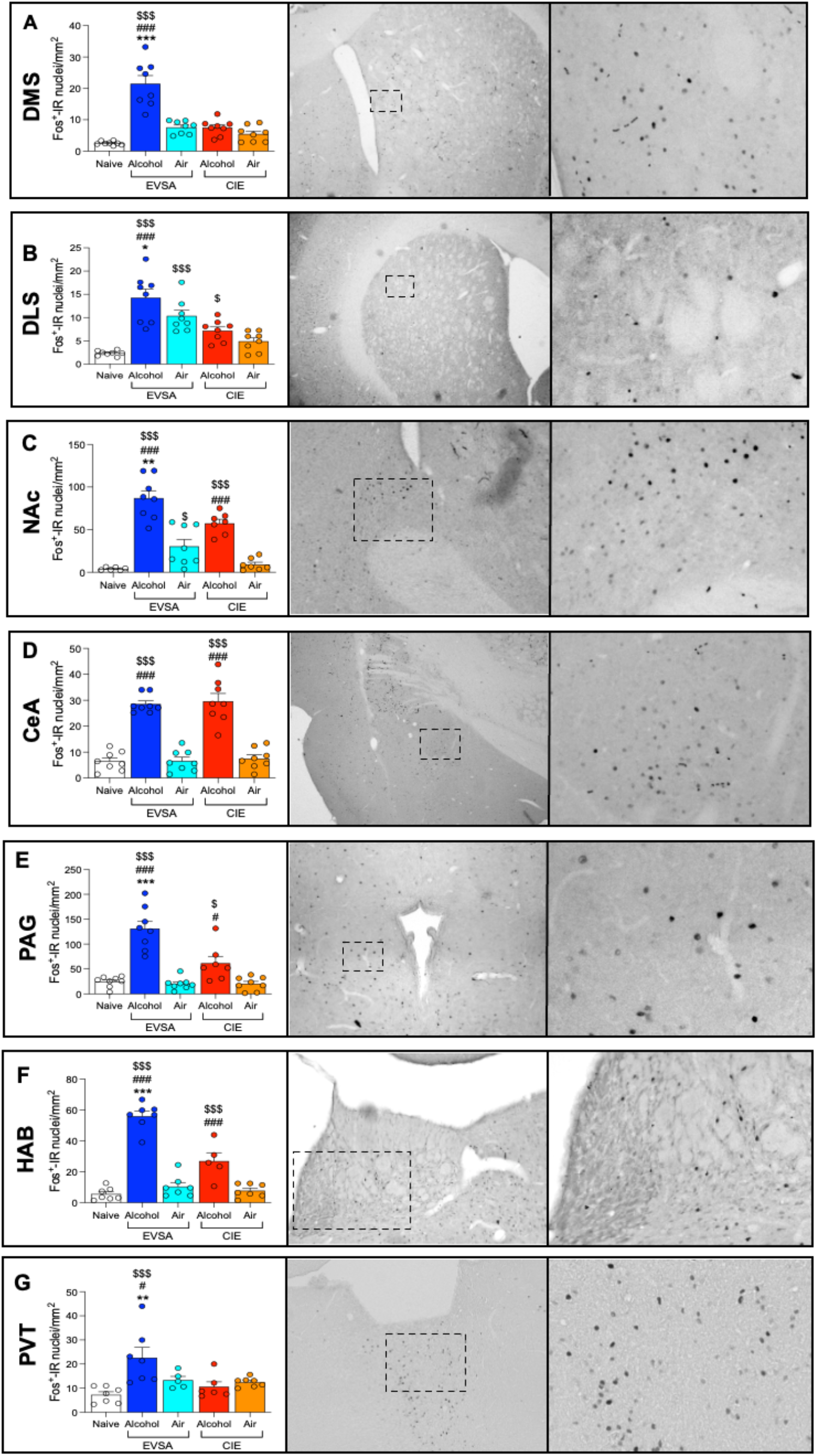
Neuronal ensembles activated by alcohol withdrawal from CIE and EVSA. Number of Fos-immunoreactive nuclei per millimeter squared and representative images of **A**) DMS, **B**) DLS, **C**) NAcc, D) CeA, E) PAG, **F**) HbL, **G**) PVT. * significant differences vs CIE, # significant differences vs air control and $ significant differences vs naïve.

#### Striatum

In the DMS, withdrawal from EVSA increased the number of Fos-positive neurons, as demonstrated by the one-way ANOVA (*F*_4,35_ = 29.92, *p* < 0.0001). Analysis of the Newman Keuls post hoc demonstrated that the EVSA rats had significantly higher levels of Fos-positive neurons compared to naïve (*p* < 0.0001), EVSA air (*p* < 0.0001) and CIE rats (*p* < 0.0001, Fig. 5A).

In the DLS, withdrawal from both EVSA and CIE increased the number of Fos-positive neurons, as demonstrated by the one-way ANOVA (*F*_4,35_ = 17.33, *p* < 0.0001). However, analysis of the Newman Keuls post hoc showed that the increase of Fos-positive cells was significantly higher in the EVSA rats compared to the CIE rats (*p* < 0.05 Fig. 5B). Interestingly, an increase in Fos-positive neurons compared to naïve animals was detected also in EVSA air control rats (*p* < 0.001 vs naïve, Fig. 5B).

#### Nucleus Accumbens

In the NAcc, withdrawal from both EVSA and CIE increased the number of Fos-positive neurons, as demonstrated by the one-way ANOVA (*F*_4,31_ = 29.50, *p* < 0.0001). However, analysis of the Newman Keuls post hoc showed that the increase of Fos-positive cells was significantly higher in the EVSA rats compared to the CIE rats (*p* < 0.01 Fig. 5C). A slight increase in Fos-positive neurons compared to naïve animals was detected also in EVSA air control rats (*p* < 0.05 vs naïve, Fig. 5C).

#### Central Amygdala

In the CeA, withdrawal from both EVSA and CIE increased the number of Fos-positive neurons, as demonstrated by the one-way ANOVA (*F*_4,35_ = 44.90, *p* < 0.0001). The Newman Keuls post hoc demonstrated that the increase in Fos positive neurons was similar during withdrawal from CIE or EVSA (*p* < 0.001 vs naïve and *p* < 0.001 vs their respective air control, Fig. 5D).

#### Periaqueductal Grey

In the PAG, withdrawal from both EVSA and CIE increased the number of Fos-positive neurons, as demonstrated by the one-way ANOVA (*F*_4,34_ = 26.12, *p* < 0.0001). However, analysis of the Newman Keuls post hoc showed that the increase of Fos-positive cells was significantly higher in the EVSA rats compared to the CIE rats (*p* < 0.001 Fig. 5E).

#### Medial Habenula

In the MHbl, withdrawal from both EVSA and CIE increased the number of Fos-positive neurons, as demonstrated by the one-way ANOVA (*F*_4,31_ = 29.50, *p* < 0.0001). However, analysis of the Newman Keuls post hoc showed that the increase of Fos-positive cells was significantly higher in the EVSA rats compared to the CIE rats (*p* < 0.001 Fig. 5F).

#### Paraventricular nucleus of the Thalamus

In the PVT, withdrawal from EVSA increased the number of Fos-positive neurons, as demonstrated by the one-way ANOVA (*F*_4,35_ = 29.92, *p* < 0.0001). Analysis of the Newman Keuls post hoc demonstrated that the EVSA rats had significantly higher levels of Fos-positive neurons compared to naïve (*p* < 0.001), EVSA air (*p* < 0.05) and CIE rats (*p* < 0.01, Fig. 5G).

## Discussion

This study demonstrates that CIE and EVSA rats exhibit similar BAL (150-400 mg% range) and similar escalation of alcohol drinking during withdrawal, while no changes in terms of drinking were observed in the air exposed rats. CIE and EVSA also increased the motivation for alcohol compared to their respective air control groups under a progressive ratio schedule of reinforcement. Acute withdrawal from EVSA and CIE recruited a similar number of Fos+ neurons in the CeA, however, acute withdrawal from EVSA recruited a higher number of Fos+ neurons in every other brain region analyzed compared to acute withdrawal from CIE. Moreover, acute withdrawal from EVSA specifically recruited the DMS and PVT, a pattern not observed in CIE rats.

The present study is a follow up of our previous work that characterized a novel animal model animal for the induction and maintenance of alcohol dependence in outbred rats using chronic intermittent alcohol vapor self-administration [39]. In our previous work we demonstrated that rats can voluntarily self-administer alcohol vapor to the point of becoming dependent on alcohol including somatic signs, anxiety-like behavior and hyperalgesia. However, it is unknown whether EVSA leads to an escalation of alcohol drinking *per se*, and whether such escalation is associated with neuroadaptations in brain regions related to stress, reward, and habit. In this new set of experiments, we wanted to evaluate if animals exposed to this new paradigm of EVSA would also drink more alcohol during acute withdrawal from vapor. To test this hypothesis, we compared the levels of alcohol drinking during withdrawal in animals passively exposed to alcohol (CIE) and animals subjected to the new vapor self-administration paradigm (EVSA). The results showed similar blood alcohol levels between CIE and EVSA rats at the end of the vapor exposure. EVSA and CIE rats similarly escalated their alcohol intake during acute withdrawal to alcohol vapor and similarly showed increased motivation for alcohol when compared to their air control counterparts. These results demonstrate that rats in the EVSA model, not only escalated their exposure to ethanol vapor, but also escalated their alcohol drinking in an operant chamber and showed increased motivation for alcohol that was similar to CIE rats. The levels of alcohol drinking after EVSA were similar to the levels described in the literature with the CIE model and relevant for alcohol dependence [12,22,23,33,40].

We next performed neuroanatomical analysis of the immediate early gene *c-fos* in several addiction-related brain regions and compared patterns of activation between CIE and EVSA rats during acute alcohol withdrawal (16 h), at the time the animals were supposed to be tested for their alcohol drinking levels. Particularly, we focused on the striatum, the nucleus accumbens, the central amygdala, the periaqueductal grey, the medial habenula and the paraventricular nucleus of the thalamus, key regions for stress, reward, and habit-related processes.

### Striatum

The results showed that acute withdrawal recruits neurons in DMS selectively in the EVSA rats. In the DLS instead, acute withdrawal recruits neurons in all the groups except for the CIE air controls, when compared to naïve. However, EVSA rats showed a significantly higher number of Fos positive cells than CIE rats, despite reaching the same BALs (Fig 4A). These results suggest that hyperactivation of the DMS and DLS in EVSA may contribute to voluntary maintenance of alcohol dependence. Indeed, the striatum is the major input station of the basal ganglia known to control goal-directed and habit-like behaviors. The striatum receives glutamatergic inputs from the prefrontal cortex and to a lesser degree from the thalamus, as well as dopaminergic inputs from the ventral midbrain, and serotonergic inputs from the dorsal raphe nucleus and glutamatergic and GABAergic input from the amygdala. Alcohol alters the function of striatal circuits in multiple ways, which may contribute to acute intoxication, alcohol seeking, dependence and withdrawal. Growing evidence indicates regional-specificity of alcohol actions on the striatum [41–43]. The dorsolateral striatum (DLS) contributes to habit formation and may participate in the development of habitual alcohol use while the dorsomedial striatum (DMS) participates in control of goal-directed actions [44], and thus may influence goal-directed alcohol seeking. The selective increased in Fos+ neurons in the DMS in EVSA rats is likely due to the fact that EVSA rats had to perform an operant response to obtain alcohol vapor while CIE rats received alcohol vapor passively. These results suggest that the EVSA model may be more useful than the CIE model to study alcohol-induced neuroadaptations in the striatum that are relevant to the increased motivation to seek alcohol in dependent subjects.

### Nucleus Accumbens and CeA

We observed increased activation of the nucleus accumbens and the central amygdala in both CIE and EVSA rats during acute alcohol withdrawal, compared to air controls and naïve, suggesting that the mechanism behind the dysregulation of these two brain regions may be similar in rats made dependent with these two models (albeit the number of Fos+ neurons in the nucleus accumbens was slighlty higher in the EVSA rats, compared to CIE animals). Converging lines of evidence show that alcohol acts on specific elements of the ventral forebrain, such as the NAcc and CeA to produce its acute positive reinforcing effects [45] and that the transition to dependence and addiction is believed to manifest as a function of dysregulated reward and stress circuitry within the the NAcc and CeA [46]. Numerous studies have identified a key role for the central nucleus of the amygdala (CeA) in alcohol drinking and alcohol dependence [18,23,47–51]. Chronic alcohol use alters CeA neuronal transmission [24,52–55], and the CeA has been shown to encode alcohol-related memories [56]. Moreover, we have shown that the activation of a specific neuronal ensemble in the CeA during alcohol withdrawal is associated with high levels of alcohol drinking in alcohol-dependent rats [18]. The similar level of Fos activation on the CeA between EVSA and CIE rats is in line with the similar BALs and escalation of alcohol drinking in both groups, suggesting that activation of the CeA may be related to the motivation to drink alcohol during acute withdrawal [45,57], but not whether the maintenance of alcohol dependence is volitional or not.

### Periaqueductal Gray

Acute withdrawal produced a robust activation that was twice as strong in EVSA comparted to CIE rats. The PAG is a major output of the extended amygdala and orchestrates emotional expression. Particularly relevant to negative emotional state, alcohol withdrawal-induced hyperalgesia is mediated in part by amygdala (CeA) projections to the PAG [58]. Glutamatergic projections from the PAG to the ventral tegmental area synapse on both GABA and dopamine neurons [59] and provide a circuit for drug seeking via the PAG. Interestingly, the PAG is also involved in the regulation of goal-directed behavior [60], particularly in the presence of an aversive stimulus [60]. Thus, PAG circuitry mediates negative emotional states and may close off the motivational loop that links negative reinforcement to drug seeking and the increased Fos activation in EVSA rats may contribute to the volitional maintenance of alcohol dependence of EVSA compared to CIE rats.

### Medial Habenula

Acute withdrawal recruited the Medial Habenula (MHb) in both EVSA and CIE rats, with ~3x higher Fos+ cells in the EVSA rats. The MHb has been shown to be involved in addiction, anxiety, and mood regulation [61,62]. It has been postulated that the LHb might be more implicated in the initial stages of recreational drug intake (associated with positive reinforcement), while the gradual shift towards compulsive drug use in addiction, strongly associated with negative affect (negative reinforcement), might be mediated a gradually greater involvement of the MHb [61,63]. Importantly, we have recently shown MHb regulation of excessive alcohol drinking in dependent rats with high addiction-like behaviors [29], demonstrating that the MHb involvement could be recruited during alcohol dependence. One possibility is that EVSA rats may develop higher negative emotional states during alcohol withdrawal, that could explain the higher activation observed in the MHb. However, in the current study we only compared CIE and EVSA rats for their levels of drinking, motivation for alcohol and BALs, but we did not look at other aspects such as somatic or emotional (anxiety and hyperalgesia) withdrawal signs. Such follow up studies will be important to test this hypothesis.

### Paraventricular Nucleus of the Thalamus

Acute withdrawal recruited the PVT only in EVSA rats. The paraventricular nucleus of the thalamus (PVT) is a midline thalamic brain region that has emerged as a critical circuit node in the regulation of behaviors across domains of affect and motivation, stress responses, and alcohol- and drug-related behaviors [64–66]. Recent studies have shown that the PVT is involved in ethanol drinking [67,68], but acute abstinence from CIE did not increase Fos activation in the PVT in mice [69]. This is in line with our data that show selective activation of the PVT in the EVSA rats, demonstrating that hyperactivation of the PVT might contribute to the voluntary maintenance of alcohol dependence in rats.

In summary, these results demonstrated that rats made dependent through the EVSA model showed phenotypical characteristic (escalation of drinking, increased motivation for alcohol, high BALs) that were similar to animals made dependent with the CIE model. However, the EVSA rats showed higher level of Fos activation in the DLS, NAcc, PAG, and HbL compared to CIE rats and showed selective recruitment of neuronal ensembles in the DMS and PVT. These results further validate the relevance of the EVSA model to the initiation and maintenance of alcohol dependence and demonstrate that escalation of alcohol vapor is associated with escalation of alcohol drinking and the selective recruitment of neuronal ensembles in brain regions critical for reward, stress, and habit-related processes. Targeting specifically these ensembles in the DMS and the PVT during acute abstinence from ethanal vapor self-administration may shed lights on the mechanisms underlying the volitional initiation and maintenance of alcohol dependence.

## Author Contributions

GdG and OG designed the research. GdG, MK, DK, performed the behavioral experiments. SS, AK, MC and RB performed analyzed the tissue samples. GdG analyzed the data and prepared the manuscript. OG edited the manuscript. All the authors reviewed the final version of the manuscript.

## References

1 Rodd ZA, Bell RL, Sable HJ, Murphy JM, McBride WJ. Recent advances in animal models of alcohol craving and relapse. Pharmacol Biochem Behav. 2004;79(3):439–50.

2 Lieber CS, DeCarli LM. The feeding of alcohol in liquid diets: two decades of applications and 1982 update. Alcohol Clin Exp Res. 1982;6(4):523–31.

3 Gilpin NW, Smith AD, Cole M, Weiss F, Koob GF, Richardson HN. Operant behavior and alcohol levels in blood and brain of alcohol-dependent rats. Alcohol Clin Exp Res. 2009;33(12):2113–23.

4 Overstreet DH, Knapp DJ, Breese GR. Similar anxiety-like responses in male and female rats exposed to repeated withdrawals from ethanol. Pharmacol Biochem Behav. 2004;78(3):459–64.

5 Aujla H, Cannarsa R, Romualdi P, Ciccocioppo R, Martin-Fardon R, Weiss F. Modification of anxiety-like behaviors by nociceptin/orphanin FQ (N/OFQ) and time-dependent changes in N/OFQ-NOP gene expression following ethanol withdrawal. Addict Biol. 2013;18(3):467–79.

6 Sidhpura N, Weiss F, Martin-Fardon R. Effects of the mGlu2/3 agonist LY379268 and the mGlu5 antagonist MTEP on ethanol seeking and reinforcement are differentially altered in rats with a history of ethanol dependence. Biological psychiatry. 2010;67(9):804–11.

7 Braconi S, Sidhpura N, Aujla H, Martin-Fardon R, Weiss F, Ciccocioppo R. Revisiting intragastric ethanol intubation as a dependence induction method for studies of ethanol reward and motivation in rats. Alcohol Clin Exp Res. 2010;34(3):538–44.

8 de Guglielmo G, Martin-Fardon R, Teshima K, Ciccocioppo R, Weiss F. MT-7716, a potent NOP receptor agonist, preferentially reduces ethanol seeking and reinforcement in post-dependent rats. Addict Biol. 2015;200:643–51.

9 Goldstein DB, Pal N. Alcohol dependence produced in mice by inhalation of ethanol: grading the withdrawal reaction. Science. 1971;172(3980):288–90.

10 Schulteis G, Markou A, Cole M, Koob GF. Decreased brain reward produced by ethanol withdrawal. Proc Natl Acad Sci U S A. 1995;92(13):5880–4.

11 Roberts AJ, Cole M, Koob GF. Intra-amygdala muscimol decreases operant ethanol self-administration in dependent rats. Alcohol Clin Exp Res. 1996;20(7):1289–98.

12 O’Dell LE, Roberts AJ, Smith RT, Koob GF. Enhanced alcohol self-administration after intermittent versus continuous alcohol vapor exposure. Alcohol Clin Exp Res. 2004;28(11):1676–82.

13 Rimondini R, Arlinde C, Sommer W, Heilig M. Long-lasting increase in voluntary ethanol consumption and transcriptional regulation in the rat brain after intermittent exposure to alcohol. FASEB J. 2002;16(1):27–35.

14 Vendruscolo LF, Roberts AJ. Operant alcohol self-administration in dependent rats: focus on the vapor model. Alcohol. 2014;48(3):277–86.

15 Gilpin NW, Richardson HN, Cole M, Koob GF. Vapor inhalation of alcohol in rats. Curr Protoc Neurosci. 2008;Chapter 9:Unit 9 29.

16 McBride WJ, Li TK. Animal models of alcoholism: neurobiology of high alcohol-drinking behavior in rodents. Crit Rev Neurobiol. 1998;12(4):339–69.

17 Ciccocioppo R, Economidou D, Cippitelli A, Cucculelli M, Ubaldi M, Soverchia L, et al. Genetically selected Marchigian Sardinian alcohol-preferring (msP) rats: an animal model to study the neurobiology of alcoholism. Addict Biol. 2006;11(3-4):339–55.

18 de Guglielmo G, Crawford E, Kim S, Vendruscolo LF, Hope BT, Brennan M, et al. Recruitment of a Neuronal Ensemble in the Central Nucleus of the Amygdala Is Required for Alcohol Dependence. The Journal of neuroscience: the official journal of the Society for Neuroscience. 2016;36(36):9446–53.

19 Leao RM, Cruz FC, Vendruscolo LF, de Guglielmo G, Logrip ML, Planeta CS, et al. Chronic nicotine activates stress/reward-related brain regions and facilitates the transition to compulsive alcohol drinking. The Journal of neuroscience: the official journal of the Society for Neuroscience. 2015;35(15):6241–53.

20 Becker HC, Lopez MF. Increased ethanol drinking after repeated chronic ethanol exposure and withdrawal experience in C57BL/6 mice. Alcohol Clin Exp Res. 2004;28(12):1829–38.

21 Lopez MF, Becker HC. Operant ethanol self-administration in ethanol dependent mice. Alcohol. 2014;48(3):295–9.

22 Richardson HN, Lee SY, O’Dell LE, Koob GF, Rivier CL. Alcohol self-administration acutely stimulates the hypothalamic-pituitary-adrenal axis, but alcohol dependence leads to a dampened neuroendocrine state. The European journal of neuroscience. 2008;28(8):1641–53.

23 de Guglielmo G, Kallupi M, Pomrenze MB, Crawford E, Simpson S, Schweitzer P, et al. Inactivation of a CRF-dependent amygdalofugal pathway reverses addiction-like behaviors in alcohol-dependent rats. Nat Commun. 2019;10(1):1238.

24 Varodayan FP, de Guglielmo G, Logrip ML, George O, Roberto M. Alcohol Dependence Disrupts Amygdalar L-Type Voltage-Gated Calcium Channel Mechanisms. The Journal of neuroscience: the official journal of the Society for Neuroscience. 2017;37(17):4593–603.

25 Kallupi M, Vendruscolo LF, Carmichael CY, George O, Koob GF, Gilpin NW. Neuropeptide YY(2)R blockade in the central amygdala reduces anxiety-like behavior but not alcohol drinking in alcohol-dependent rats. Addict Biol. 2014;19(5):755–7.

26 Vendruscolo LF, Barbier E, Schlosburg JE, Misra KK, Whitfield TW, Jr., Logrip ML, et al. Corticosteroid-dependent plasticity mediates compulsive alcohol drinking in rats. J Neurosci. 2012;32(22):7563–71.

27 Vendruscolo LF, Estey D, Goodell V, Macshane LG, Logrip ML, Schlosburg JE, et al. Glucocorticoid receptor antagonism decreases alcohol seeking in alcohol-dependent individuals. J Clin Invest. 2015;125(8):3193–7.

28 Kimbrough A, de Guglielmo G, Kononoff J, Kallupi M, Zorrilla EP, George O. CRF1 Receptor-Dependent Increases in Irritability-Like Behavior During Abstinence from Chronic Intermittent Ethanol Vapor Exposure. Alcohol Clin Exp Res. 2017;41(11):1886–95.

29 Kononoff J, Kallupi M, Kimbrough A, Conlisk D, de Guglielmo G, George O. Systemic and Intra-Habenular Activation of the Orphan G Protein-Coupled Receptor GPR139 Decreases Compulsive-Like Alcohol Drinking and Hyperalgesia in Alcohol-Dependent Rats. eNeuro. 2018;5(3).

30 Funk CK, O’Dell LE, Crawford EF, Koob GF. Corticotropin-releasing factor within the central nucleus of the amygdala mediates enhanced ethanol self-administration in withdrawn, ethanol-dependent rats. The Journal of neuroscience: the official journal of the Society for Neuroscience. 2006;26(44):11324–32.

31 Healey JC, Winder DG, Kash TL. Chronic ethanol exposure leads to divergent control of dopaminergic synapses in distinct target regions. Alcohol. 2008;42(3):179–90.

32 Nealey KA, Smith AW, Davis SM, Smith DG, Walker BM. kappa-opioid receptors are implicated in the increased potency of intra-accumbens nalmefene in ethanol-dependent rats. Neuropharmacology. 2011;61(1-2):35–42.

33 Smith AW, Nealey KA, Wright JW, Walker BM. Plasticity associated with escalated operant ethanol self-administration during acute withdrawal in ethanol-dependent rats requires intact matrix metalloproteinase systems. Neurobiol Learn Mem. 2011;96(2):199–206.

34 Valdez GR, Roberts AJ, Chan K, Davis H, Brennan M, Zorrilla EP, et al. Increased ethanol self-administration and anxiety-like behavior during acute ethanol withdrawal and protracted abstinence: regulation by corticotropin-releasing factor. Alcohol Clin Exp Res. 2002;26(10):1494–501.

35 Walker BM. Conceptualizing withdrawal-induced escalation of alcohol self-administration as a learned, plasticity-dependent process. Alcohol. 2012;46(4):339–48.

36 Walker BM, Koob GF. Pharmacological evidence for a motivational role of kappa-opioid systems in ethanol dependence. Neuropsychopharmacology. 2008;33(3):643–52.

37 Walker BM, Zorrilla EP, Koob GF. Systemic kappa-opioid receptor antagonism by nor-binaltorphimine reduces dependence-induced excessive alcohol self-administration in rats. Addict Biol. 2011;16(1):116–9.

38 de Guglielmo G, Conlisk DE, Barkley-Levenson AM, Palmer AA, George O. Inhibition of Glyoxalase 1 reduces alcohol self-administration in dependent and nondependent rats. Pharmacol Biochem Behav. 2018;167:36–41.

39 de Guglielmo G, Kallupi M, Cole MD, George O. Voluntary induction and maintenance of alcohol dependence in rats using alcohol vapor self-administration. Psychopharmacology (Berl). 2017.

40 Kissler JL, Walker BM. Dissociating Motivational From Physiological Withdrawal in Alcohol Dependence: Role of Central Amygdala kappa-Opioid Receptors. Neuropsychopharmacology. 2016;41(2):560–7.

41 Corbit LH, Nie H, Janak PH. Habitual alcohol seeking: time course and the contribution of subregions of the dorsal striatum. Biological psychiatry. 2012;72(5):389–95.

42 Jeanblanc J, He DY, Carnicella S, Kharazia V, Janak PH, Ron D. Endogenous BDNF in the dorsolateral striatum gates alcohol drinking. The Journal of neuroscience: the official journal of the Society for Neuroscience. 2009;29(43):13494–502.

43 Chen G, Cuzon Carlson VC, Wang J, Beck A, Heinz A, Ron D, et al. Striatal involvement in human alcoholism and alcohol consumption, and withdrawal in animal models. Alcohol Clin Exp Res. 2011;35(10):1739–48.

44 Balleine BW, O’Doherty JP. Human and rodent homologies in action control: corticostriatal determinants of goal-directed and habitual action. Neuropsychopharmacology. 2010;35(1):48–69.

45 Koob GF. Alcoholism: allostasis and beyond. Alcohol Clin Exp Res. 2003;27(2):232–43.

46 Koob GF. Addiction is a Reward Deficit and Stress Surfeit Disorder. Front Psychiatry. 2013;4:72.

47 Gilpin NW, Herman MA, Roberto M. The central amygdala as an integrative hub for anxiety and alcohol use disorders. Biological psychiatry. 2015;77(10):859–69.

48 Koob GF. Brain stress systems in the amygdala and addiction. Brain Res. 2009;1293:61–75.

49 Koob GF, Volkow ND. Neurocircuitry of addiction. Neuropsychopharmacology. 2010;35(1):217–38.

50 Wrase J, Makris N, Braus DF, Mann K, Smolka MN, Kennedy DN, et al. Amygdala volume associated with alcohol abuse relapse and craving. Am J Psychiatry. 2008;165(9):1179–84.

51 Kimbrough A, Lurie DJ, Collazo A, Kreifeldt M, Sidhu H, Macedo GC, et al. Brain-wide functional architecture remodeling by alcohol dependence and abstinence. Proc Natl Acad Sci U S A. 2020;117(4):2149–59.

52 Roberto M, Gilpin NW, Siggins GR. The central amygdala and alcohol: role of gammaaminobutyric acid, glutamate, and neuropeptides. Cold Spring Harb Perspect Med. 2012;2(12):a012195.

53 Roberto M, Madamba SG, Moore SD, Tallent MK, Siggins GR. Ethanol increases GABAergic transmission at both pre- and postsynaptic sites in rat central amygdala neurons. Proc Natl Acad Sci U S A. 2003;100(4):2053–8.

54 Roberto M, Madamba SG, Stouffer DG, Parsons LH, Siggins GR. Increased GABA release in the central amygdala of ethanol-dependent rats. The Journal of neuroscience: the official journal of the Society for Neuroscience. 2004;24(45):10159–66.

55 Roberto M, Schweitzer P, Madamba SG, Stouffer DG, Parsons LH, Siggins GR. Acute and chronic ethanol alter glutamatergic transmission in rat central amygdala: an in vitro and in vivo analysis. The Journal of neuroscience: the official journal of the Society for Neuroscience. 2004;24(7):1594–603.

56 Barak S, Liu F, Ben Hamida S, Yowell QV, Neasta J, Kharazia V, et al. Disruption of alcohol-related memories by mTORC1 inhibition prevents relapse. Nat Neurosci. 2013;16(8):1111–7.

57 Koob GF, Le Moal M. Addiction and the brain antireward system. Annu Rev Psychol. 2008;59:29–53.

58 Avegno EM, Lobell TD, Itoga CA, Baynes BB, Whitaker AM, Weera MM, et al. Central Amygdala Circuits Mediate Hyperalgesia in Alcohol-Dependent Rats. The Journal of neuroscience: the official journal of the Society for Neuroscience. 2018;38(36):7761–73.

59 Omelchenko N, Sesack SR. Periaqueductal gray afferents synapse onto dopamine and GABA neurons in the rat ventral tegmental area. J Neurosci Res. 2010;88(5):981–91.

60 Mobbs D, Petrovic P, Marchant JL, Hassabis D, Weiskopf N, Seymour B, et al. When fear is near: threat imminence elicits prefrontal-periaqueductal gray shifts in humans. Science. 2007;317(5841):1079–83.

61 Batalla A, Homberg JR, Lipina TV, Sescousse G, Luijten M, Ivanova SA, et al. The role of the habenula in the transition from reward to misery in substance use and mood disorders. Neurosci Biobehav Rev. 2017;80:276–85.

62 Fowler CD, Kenny PJ. Habenular signaling in nicotine reinforcement. Neuropsychopharmacology. 2012;37(1):306–7.

63 Loonen AJM, Kupka RW, Ivanova SA. Circuits Regulating Pleasure and Happiness in Bipolar Disorder. Front Neural Circuits. 2017;11:35.

64 Hartmann MC, Pleil KE. Circuit and neuropeptide mechanisms of the paraventricular thalamus across stages of alcohol and drug use. Neuropharmacology. 2021;198:108748.

65 Matzeu A, Martin-Fardon R. Drug Seeking and Relapse: New Evidence of a Role for Orexin and Dynorphin Co-transmission in the Paraventricular Nucleus of the Thalamus. Front Neurol. 2018;9:720.

66 Matzeu A, Martin-Fardon R. Blockade of Orexin Receptors in the Posterior Paraventricular Nucleus of the Thalamus Prevents Stress-Induced Reinstatement of Reward-Seeking Behavior in Rats With a History of Ethanol Dependence. Front Integr Neurosci. 2020;14:599710.

67 Barson JR, Ho HT, Leibowitz SF. Anterior thalamic paraventricular nucleus is involved in intermittent access ethanol drinking: role of orexin receptor 2. Addict Biol. 2015;20(3):469–81.

68 Barson JR, Poon K, Ho HT, Alam MI, Sanzalone L, Leibowitz SF. Substance P in the anterior thalamic paraventricular nucleus: promotion of ethanol drinking in response to orexin from the hypothalamus. Addict Biol. 2017;22(1):58–69.

69 Smith RJ, Anderson RI, Haun HL, Mulholland PJ, Griffin WC, 3rd, Lopez MF, et al. Dynamic c-Fos changes in mouse brain during acute and protracted withdrawal from chronic intermittent ethanol exposure and relapse drinking. Addict Biol. 2020;25(6):e12804.

